# Bayesian Inference of Chemical Mixtures in Risk Assessment Incorporating the Hierarchical Principle

**DOI:** 10.1101/2023.05.19.541480

**Authors:** Debamita Kundu, SungDuk Kim, Paul S. Albert

**Affiliations:** Biostatistics Division, University of Virginia, VA, USA; Biostatistics Branch, National cancer Institute, MD, USA

**Keywords:** Chemical mixture, Interaction, Shrinkage, Collapsed Gibbs

## Abstract

Analyzing health effects associated with exposure to environmental chemical mixtures is a challenging problem in epidemiology, toxicology, and exposure science. In particular, when there are a large number of chemicals under consideration it is difficult to estimate the interactive effects without incorporating reasonable prior information. Based on substantive considerations, researchers believe that true interactions between chemicals need to incorporate their corresponding main effects. In this paper, we use this prior knowledge through a shrinkage prior that a *priori* assumes an interaction term can only occur when the corresponding main effects exist. Our initial development is for logistic regression with linear chemical effects. We extend this formulation to include non-linear exposure effects and to account for exposure subject to detection limit. We develop an MCMC algorithm using a shrinkage prior that shrinks the interaction terms closer to zero as the main effects get closer to zero. We examine the performance of our methodology through simulation studies and illustrate an analysis of chemical interactions in a case-control study in cancer.

## 1 Introduction

Assessing the health effects of environmental chemical mixtures is an important challenge in environmental epidemiology. It is quite difficult to find the relationship between exposure variables and a health outcome when the effects may be non-linear, interactions effects are present, and may be subject to detection limits. In the past decade, there have been numerous approaches for analyzing this type of data. Hwang et al. (2019); Zhang et al. (2012) proposed latent class approach that links the exposure profile with disease severity with latent variables. Bobb et al. (2015) employed Bayesian kernal machine regression that allows for flexible non-linear estimation of chemical effects on disease outcomes. Further, Carrico et al. (2015) proposed a weighted quantile sum approach that generalizes using the cumulative chemical exposure for predicting disease outcomes. Herring (2010) proposed a Bayesian regression approach that estimates linear main and interactive effects using a Dirichlet process prior and incorporates detection limits for the chemical exposures.

In this paper, we propose a Bayesian approach that incorporates non-linear exposure, interaction effects, and detection limits in a flexible way that does not require parametric assumptions on the exposure distributions. When the number of parameters are moderately large, the inclusion of all pairwise interaction terms results in sparsity that in turn results in poor estimation with maximum-likelihood estimation. We propose a shrinkage prior approach for estimating interactions. First, we investigate a shrinkage prior where interactions are treated the same as main effects and no relationship between the two is incorporated. In a second approach, we propose a shrinkage prior that incorporates a relationship between the interaction and main effects to increase performance in sparse data situations. The hierarchical principle in linear models specifies that interactions will only be examined in situations where the corresponding main effects are sizable (McCullagh and Nelder, 1989). Models with interactions without main effects place restrictions on the parameter space that are not natural. We will show how incorporating this additional structure will provide efficiency gain when studying the interactions in chemical mixtures. Additionally, we show how to extend the shrinkage prior approach to situations where multiple parameters are used to model the effect of each chemical component on disease risk. This later extension can be used to model non-linear exposure effects as well as to provide a flexible approach for dealing with detection limits in mixtures. We propose a shrinkage prior approach that incorporates the hierarchical principle for estimation of interactions for linear and nonlinear exposures as well as for exposures subject to detection limits.

In our paper, we describe our methodology in detail in Section 2. Next in Section 3, we discuss our prior specifications and posterior computations. In Section 4 we describe how to extend our model from linear where the relationship of an exposure has a single parameter to the situation where multiple parameters are associated with that exposure (e.g., nonlinear exposures or detection limits). In Section 5 we show the efficiency of our proposed method by simulation studies and results. Finally, we applied our methodology to the NCI-SEER NHL study and describe our findings in Section 6.

## 2 Model

Let *Y* = (*Y*_1_, *Y*_2_⋯, *Y*_*N*_)′ denotes the binary health response for *N* individuals and **X**_**i**_ = (*X*_*i*1_, *X*_*i*2_… ., *X*_*ip*_)′ be the corresponding *p*-dimensional vector of continuous chemical exposures. For *p* chemicals we have *p*(*p* −1)/2 two-way interactions. We consider a logistic regression model with linear effects in their corresponding interactions of the following form:

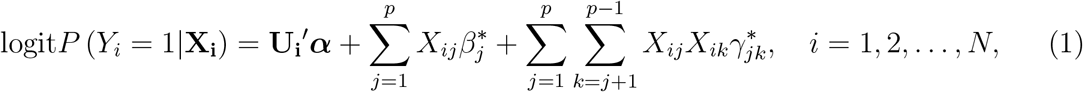

where logit 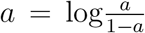 **U**_**i**_ denotes *q*-dimensional covariate vector which includes an intercept, ***α*** is the corresponding regression coefficient vector,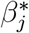 denotes the main effect regression coefficient of the *j*^*th*^ chemical, and 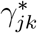 denotes the interaction effect regression coefficient of the *j*^*th*^ and *k*^*th*^ chemicals. We consider a latent variable approach (Albert and Chib, 1993) and approximate equation (1) using a robit link (Liu, 2004). Let consider ***ω*** = (*ω*_1_, *ω*_2_, …, *ω*_*N*_)′ be *N* -dimensional latent vector such that

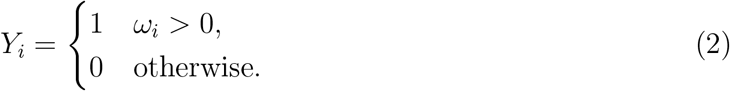

Where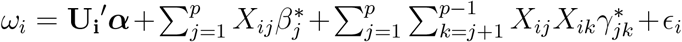 If 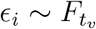 where 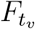 is a cumulative distribution function of student *t*-distribution with *v* degrees of freedom, it is called *robi*i*t* (*v*) regression (Lange et al., 1989), i.e. 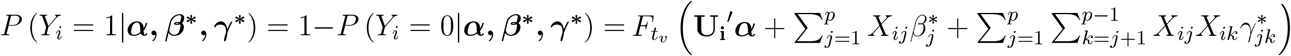 where 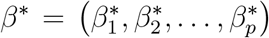 and 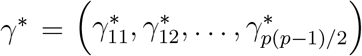

As *v→*∞ the *robit* (*v*) model becomes the probit regression model. Liu (2004) suggested that the *robit* link with *v* = 7 degrees of freedom closely approximates the *logit* link with 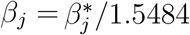 and 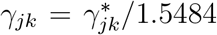 Moreover, we use the fact that the *t*-distribution can be represented as a scale mixture of normal distribution by introducing a mixing variable *λ*_*i*_, such that 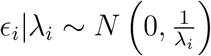 and 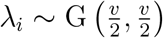 where *N* (*μ, σ*^2^) denotes a normal distribution with mean *μ* and variance *σ*^2^ and G(*c*_1_, *c*_2_) denotes the gamma distribution with mean *c*_1_*/c*_2_ and variance 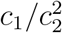 to formulate the likelihood. For simplicity, let consider for the *i*^*th*^ individual the interaction term between two exposure variables *X*_*ij*_ and *X*_*ik*_ is defined by 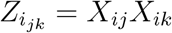 and 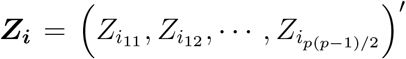Hence 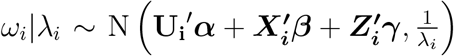 and 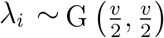 where *β* = (*β*_1_, *β*_2_, …, *β*_*p*_) and *γ* = *γ*_11_, *γ*_12_, …, *γp*(*p*−1)/_2_. From equations (1) and (2) the complete data likelihood can be written as:

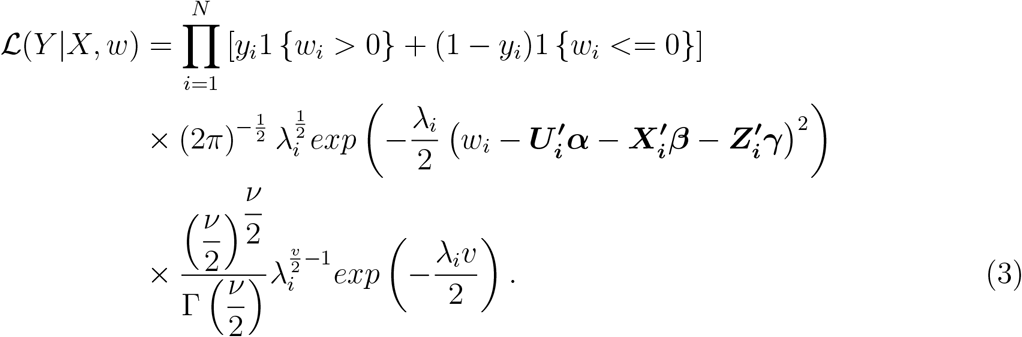

## 3 Prior & Posterior Distribution

For linear models with interactions, the hierarchical principle implies that interactions should only be included if the corresponding main effects are non-zero (Chipman et al., 1997; Griffin et al., 2017). Hence, we consider a dependence structure between the main and interaction effects such that the inclusion of interaction effects depends on the inclusion of the corresponding main effects. To this end, we consider the following prior distributions :

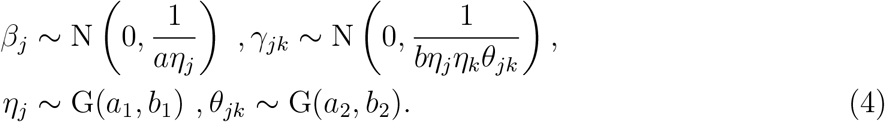

In the prior distribution in equation (4), *a* controls the global shrinkage towards the origin for the main effect regression coefficient *β*_*j*_, *j* = 1, 2, …, *p*, and ***η*** = (*η*_1_, *η*_2_, …, *η*_*p*_)′ is the predictor specific local shrinkage parameter that allow deviations in the degree of shrinkage between predictors. *a*_1_, *a*_2_ defines the shape parameter and *b*_1_, *b*_2_ defines the scale parameter of the gamma distributions. We consider a G(*a*_1_ = 1, *b*_1_ = 1) distribution as a prior choice for *η*_*j*_ with mean and variance 1 to induce some variability between the *η*_*j*_’s. In this formulation, larger values of *η*_*j*_’s will induce more shrinkage towards zero, while smaller values of *η*_*j*_ result in minimum shrinkage to zero. This specification is based on global-local shrinkage framework (Polson and Scott, 2010), where typical recommendation is to consider a heavy tail distribution for the local shrinkage parameter to avoid over-shrinking large signals, and the global shrinkage parameter should have substantial mass near zero. Similarly, for the interaction effect regression coefficients *γ*_*jk*_, the global shrinkage parameter *b* shrinks all parameter towards zero. In contrast, the predictor specific local shrinkage parameter ***θ*** = (*θ*_1_, *θ*_2_, …, *θ*_*p*(*p*−1)/2_)′ captures the interaction specific shrinkage effects. We consider a G(*a*_2_ = 1, *b*_2_ = 1) prior for *θ*_*jk*_. Note that, to share the information between main and interaction effects the prior variance of *γ*_*jk*_ is also dominated by the term *η*_*j*_*η*_*k*_. The parameter *γ*_*jk*_ is shrunk to zero if at least one of *η*_*j*_, *η*_*k*_ or *θ*_*jk*_ is large. Consistent with the hierarchical principle, an interaction term will tend to be small if either *β*_*j*_ or *β*_*k*_ is small or *θ*_*jk*_ is large. Furthermore, we also consider a G(1, 1) prior on both global shrinkage parameter *a* and *b*, and a vague prior N(0, 10^2^) for the regression coefficient *α*_*j*_. The posterior distribution based on the complete data is given by

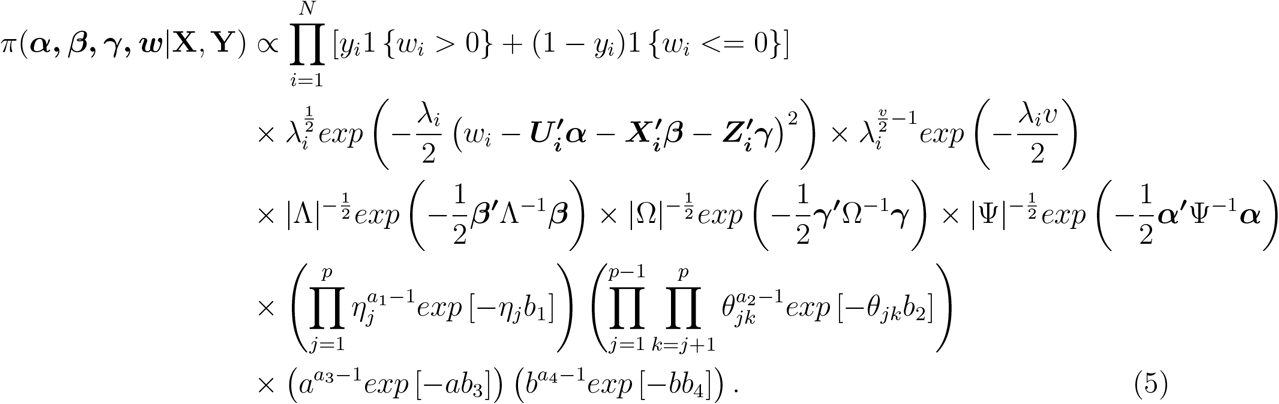

In equation (5), we de fine ***β*** = (*β* _1_, *β*_2_, …, *β*_*p*_)′, ***γ*** = (*γ*_12_, *γ*_13_, …, *γ*_*p*(*p*−1)_) ′, Λ = diag(1/*aη*_1_, 1/*aη*_2_, …, 1/*aη*_*p*_) Ω = diag (1/*bη*_1_*η*_2_*θ*_12_, 1/*bη*_1_*η*_3_*θ*_13_, …, 1/*bη*_p−1_*η*_*p*_*θ*_(*p*−1)*p*_) and Ψ= 10^2^*I*_*q*_, where *I*_*q*_ represents the *q* × *q* order identity matrix.

The proposed methodology can be easily extended to incorporate other prior distributions such as horseshoe, Cauchy, and Dirichlet-Laplace prior.

### 3.1 Computational Development

We present a detailed development of the Markov chain Monte Carlo (MCMC) sampling algorithm. The conditional posterior distributions are derived from the equation (5) and inference is performed through MCMC methods. We define 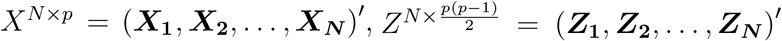. Based on the joint posterior distribution, we derive the full conditional posterior distribution of all parameters and thus get the Gibbs samples by iterating the following sampling steps:

- We sample *w*_*i*_ from the conditional posterior distribution *ω*_*i*_ |*λ*_*i*_, ***β, γ, η, θ***, *a, b*, which is a truncated normal distribution with mean 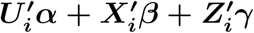 variance 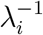 for all *i* = 1, 2, …, *N*.
- For *i* = 1, 2, …, *N*, sample *λ*_*i*_ independently from its conditional posterior distribution *λ*_*i*_ |*ω*_*i*_, ***β, γ, η, θ***, *a, b* which follows 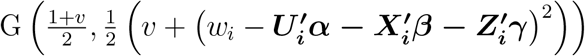
- Then, sample the covariates ***α*** from its conditional multivariate normal posterior distribution MVN (*μ*_*α*_, Σ_*α*_), where the posterior mean is ***μ***_*α*_ = (*U′*Σ^−1^*U* + Ψ^−1^)^−1^ *U′*Σ^−1^ (***w − X′β − Z′γ***), posterior variance Σ_***α***_ = (*U′*Σ^−1^*U* + Ψ^−1^)^−1^ and 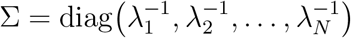
- Next we sample ***β*** from its conditional multivariate normal posterior distribution MVN (*μ*_*β*_, Σ_*β*_), where posterior mean ***μ***_***β***_ = (*X′*Σ^−1^*X* + Λ^−1^)^− 1^ *X′*Σ^−1^ (***w − U′α − Z′γ***) and posterior variance Σ_*β*_ = (*X′*Σ^−1^*X* + Λ^−1^)^− 1^
- Then, sample ***γ*** from its conditional posterior distribution MVN (*μ*_*γ*_, Σ_*γ*_), where ***μ***_*γ*_ = (*Z′*Σ^−1^*Z* + *Ω*^−1^)^−1^ *Z′*Σ^−1^ (***w − U′α − X′β***) and Σ_***γ***_ = (*Z′*Σ^−1^*Z* + *Ω*^−1^)^−1^.
- For *j* = 1, 2, …, *p*, sample *η*_*j*_ independently from its conditional posterior distribution Gamma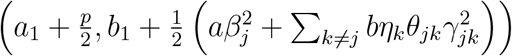
- For *j* = 1, 2, …, *p* an d *k* = *j* + 1, 2, …, *p −* 1 sample *θ*_*jk*_ from the conditional posterior distribution Gamma 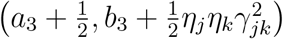
- Next, sample *a*|***β, η*** from Gamma 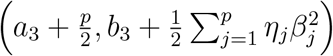
- Lastly, sample *b*|***γ, η, θ*** from Gamma 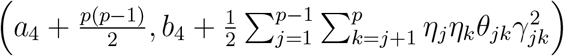

## 4 Multiple Parameters Per Exposure

Previously in Section 2, we described our method in the context of a linear exposure and outcome relationship. In many realistic settings, more than one parameter will be needed to model individual exposures in the mixtures (e.g., quadratic exposures). Hence, in this section, we extend our methodology to capture those non-linear exposure-outcome relationships using the following logistic regression model:

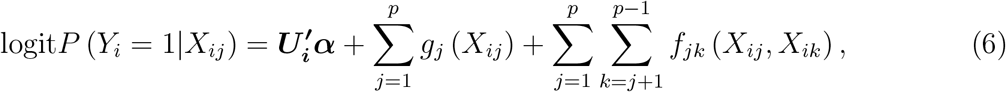

For modeling mixtures with nonlinear exposure relationships, we can use a polynomial representation for the effect of each exposure. Polynomial effects can be incorporated in the main and interaction terms by using equation (6) with functions 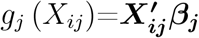 and 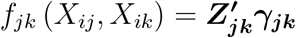 and the logistic regression can be written in the following form:

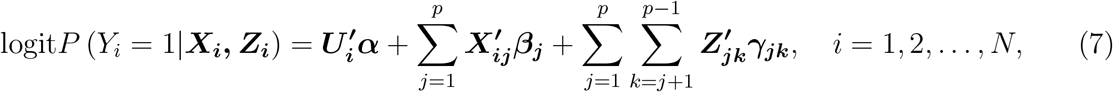

where, 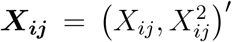 ′ and 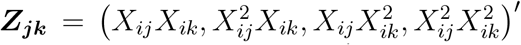 and the regression coefficients ***β***_***j***_ = (*β*_*j*1_, *β*_*j*2_)′ and ***γ***_***jk***_ = (*γ*_*jk*1_, *γ*_*jk*2_, *γ*_*jk*3_, *γ*_*jk*4_)′.

Individual components of chemical mixtures may be subject to lower detection limits where the measurements are censored below these limits. Chiou et al. (2019) and Ortega-Villa et al. (2021) proposed a two component exposure model for lower detection limits where the effect has one component indicating whether the exposure is above the detection limit and the other reflecting the value of the measurement if the exposure is above the detection limit. These authors showed that this parameterization does not make the unverifiable modelling assumptions inherent in treating lower detection limits as left censored in a parametric exposure distribution. We can incorporate detection limits into the mixture analysis by using equation (6) as:where,

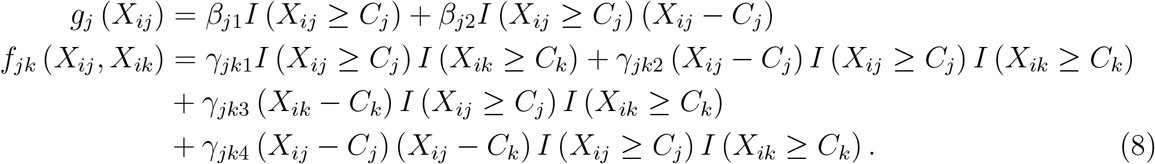

In equation (8), *β*_*j*1_ is the log odds of disease at the value of the detection limit relative to the log-odds of disease below the detection limit. Further, *β*_*j*2_ is the log-odds ratio of disease for a one unit change in exposure above the detection limit. The parameter vector *γ*_*jk*._ measure the interactive effects between the *j*^*th*^ and *k*^*th*^ chemical. Specifically, *γ*_*jk*1_ measures the interactive effect of being above the detection limit on both the *j*^*th*^ and *k*^*th*^ chemical, *γ*_*jk*4_ measures the interactive effect of increasing *X*_*ij*_ and/or *X*_*ik*_ when both markers are above the detection limit. Finally, *γ*_*jk*2_ and *γ*_*jk*3_ are cross product interaction effects.

The shared shrinkage prior proposed in Section 3 can be extended to the multiple parameter per exposure case using the grouped shrinkage prior (Casella et al., 2010). We show this for the non-linear formulation (7), but it applies more generally to other settings such as detection limits (8). Specifically,

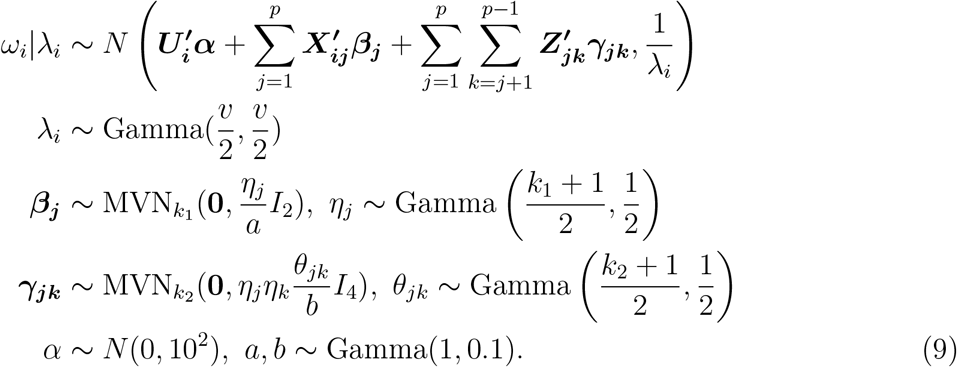

Here *k*_1_ and *k*_2_ defines the dimension of the parameters ***β***_***j***_ and ***γ***_***jk***_, respectively. In the non-linear case we share the information between main effect and interaction effect similarly as in linear hierarchical model in the equation (4). Posterior calculation follows as in Section 3.1.

## 5 Simulation study and results

We perform a series of the simulation studies to investigate the performance of the proposed methodology. We compare the proposed model with following two models.

1. **Independent Vague Prior** : We incorporate the vague prior *β*_*j*_, *γ*_*jk*_ ∼ *N* (0, 10^2^), where no dependence is introduced between the main and the interaction effects. This approach is approximately a maximum likelihood approach.
2. **Independent shrinkage prior** : Under this approach we do not share the information between main effects and interaction effects. We consider an independent shrinkage prior on both interaction effect and main effects regression coefficients. To that end, 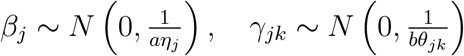 Hence in this model, the inclusion of an interaction effect is not contingent on the inclusion of main effects.

### 5.1 Simulation for linear exposure effect

We conduct a simulation study for the linear hierarchical model in equation (4), where data are generated using model in equation (2) in Section 2. For each *i*^*th*^ individual we generate *p* chemical exposures ***X***_***i***_ = (*X*_*i*1_, *X*_*i*2_, …, *X*_*ip*_)′ independently from a multivariate normal distribution with mean zero and covariance matrix *I*_*p*_. Thus we have our design matrix *X*^*N×p*^ for chemical exposure. Then we generate *p*(*p*_*−1*_)/2 interaction effects for each individual by multiplying the corresponding main effects, i.e. ***Z***_***i***_ = (*X*_*i*1_*X*_*i*2_, *X*_*i*1_*X*_*i*3_, …, *X*_*ip*_*X*_*i*(*p*−1)_) ′, where *l* = 1, 2, …, *p*(*p*_−1_)/2. Later we scaled each of the main and interaction exposures by dividing their respective standard deviations to perform the analysis. We generate 200 datasets for each model, and for each data set we run the MCMC chain for 50,000 iterations with a burn-in of 5000 iterations. We consider every 5^*th*^ sample to reduce the auto-correlation. We use those 9000 MCMC samples for posterior inferences. We consider different simulation scenarios to access the utility of our methodology in details.

- **Simulation #1:** *N* = 1000, *p* = 10 : We generate data under a model consistent with the hierarchical principle. We consider *N* = 1000 individual and 10 chemicals (*p* = 10). We further consider 5 out of 10 main effects have effect size 1 and 10 out of 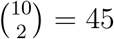interaction terms has effect size 0.5 and rest are set to zero. The intercept is set to 0.6 to have the overall prevalence rate of approximately 50%.
- **Simulation #2:** *N* = 250, *p* = 10 : In this settings, we consider a smaller sample size *N* = 250 individual and total number of chemicals *p* = 10. The main effect parameters ***β*** and interaction effect parameters ***γ*** are identical to simulation #1.
- **Simulation #3:** *N* = 1000, **Main effects and no interaction effects**: In this settings, we consider a sample size of *N* = 1000 individual and a total number of chemicals *p* = 10. We generate the data from a model with the same main effect parameters ***β*** as in simulation #1 and simulation #2, but the interaction effect parameters ***γ*** are set to zero, i.e, we have main effects but no interaction effects.
- **Simulation #4:** *N* = 1000, **Interactions with no main effects** : We generate data from a model with no main effects but with the same interaction effects as in simulation #1 and simulation #2.

#### 5.1.1 Simulation results from linear exposure effect

In this section, we compare the performance of the proposed methodology with the model with an independent vague prior and independent shrinkage model. Table 1 and 2 show the mean standard deviation and root mean square error (RMSE) for a few of the interaction terms in simulation #1 and simulation #2 respectively. First the results show that using a vague prior result in poor estimation, particularly for smaller sample size. Secondly, incorporating the shared shrinkage prior result in a large efficiency gain relative to the independent shrinkage prior. We computed the geometric mean of the relative efficiencies comparing the independent versus the shared shrinkage across all 45 interaction parameters. Shared shrinkage showed efficiency advantages in Table 1 and Table 2 with mean efficiencies of 1.25 and 1.42, respectively. As expected, the efficiency gains were larger for the smaller sample size that has increased sparsity.

**Table 1:** Results from simulation #1: *N* =1000, *p*=10

| Parameters | $\gamma$ | Independent Vague | | | Independent shrinkage | | | Shared Shrinkage | | | Relative*<br>Efficiency |
| --- | --- | --- | --- | --- | --- | --- | --- | --- | --- | --- | --- |
| | | $\hat{\gamma}$ | sd | RMSE | $\hat{\gamma}$ | sd | RMSE | $\hat{\gamma}$ | sd | RMSE | |
| Interactions | $\gamma_{12}=0.5$ | 0.892 | 0.040 | 0.472 | 0.546 | 0.023 | 0.169 | 0.533 | 0.022 | 0.151 | 1.252 |
| | $\gamma_{13}=0.5$ | 0.883 | 0.040 | 0.462 | 0.539 | 0.022 | 0.162 | 0.529 | 0.022 | 0.152 | 1.136 |
| | $\gamma_{14}=0.5$ | 0.904 | 0.041 | 0.470 | 0.553 | 0.023 | 0.154 | 0.533 | 0.022 | 0.160 | 0.925 |
| | $\gamma_{15}=0.5$ | 0.914 | 0.040 | 0.494 | 0.563 | 0.022 | 0.179 | 0.528 | 0.021 | 0.152 | 1.387 |
| | $\gamma_{16}=0$ | 0.003 | 0.024 | 0.195 | 0.001 | 0.013 | 0.113 | 0.009 | 0.011 | 0.104 | 1.179 |
| | $\gamma_{17}=0$ | -0.006 | 0.025 | 0.203 | -0.004 | 0.014 | 0.123 | 0.005 | 0.011 | 0.098 | 1.575 |
| | $\gamma_{18}=0$ | 0.005 | 0.024 | 0.184 | 0.003 | 0.013 | 0.106 | 0.009 | 0.011 | 0.091 | 1.357 |
| | $\gamma_{19}=0$ | 0.000 | 0.024 | 0.174 | -0.003 | 0.013 | 0.102 | 0.000 | 0.011 | 0.097 | 1.104 |
| | $\gamma_{1,10}=0$ | 0.003 | 0.024 | 0.199 | -0.001 | 0.013 | 0.114 | -0.005 | 0.011 | 0.103 | 1.225 |
| | $\gamma_{23}=0.5$ | 0.869 | 0.039 | 0.448 | 0.532 | 0.022 | 0.159 | 0.537 | 0.022 | 0.155 | 1.053 |
| | $\gamma_{24}=0.5$ | 0.907 | 0.041 | 0.486 | 0.559 | 0.023 | 0.175 | 0.524 | 0.021 | 0.152 | 1.325 |
| | $\gamma_{25}=0.5$ | 0.875 | 0.039 | 0.464 | 0.536 | 0.022 | 0.172 | 0.538 | 0.022 | 0.154 | 1.248 |
| | $\gamma_{26}=0$ | 0.010 | 0.024 | 0.190 | 0.005 | 0.013 | 0.110 | 0.004 | 0.011 | 0.088 | 1.563 |

\*Relative efficiency=Ratio of MSE of independent shrinkage prior and the MSE of shared shrinkage prior.
The geometric mean efficiency gain across all 45 interaction terms is 1.24.

**Table 2:**
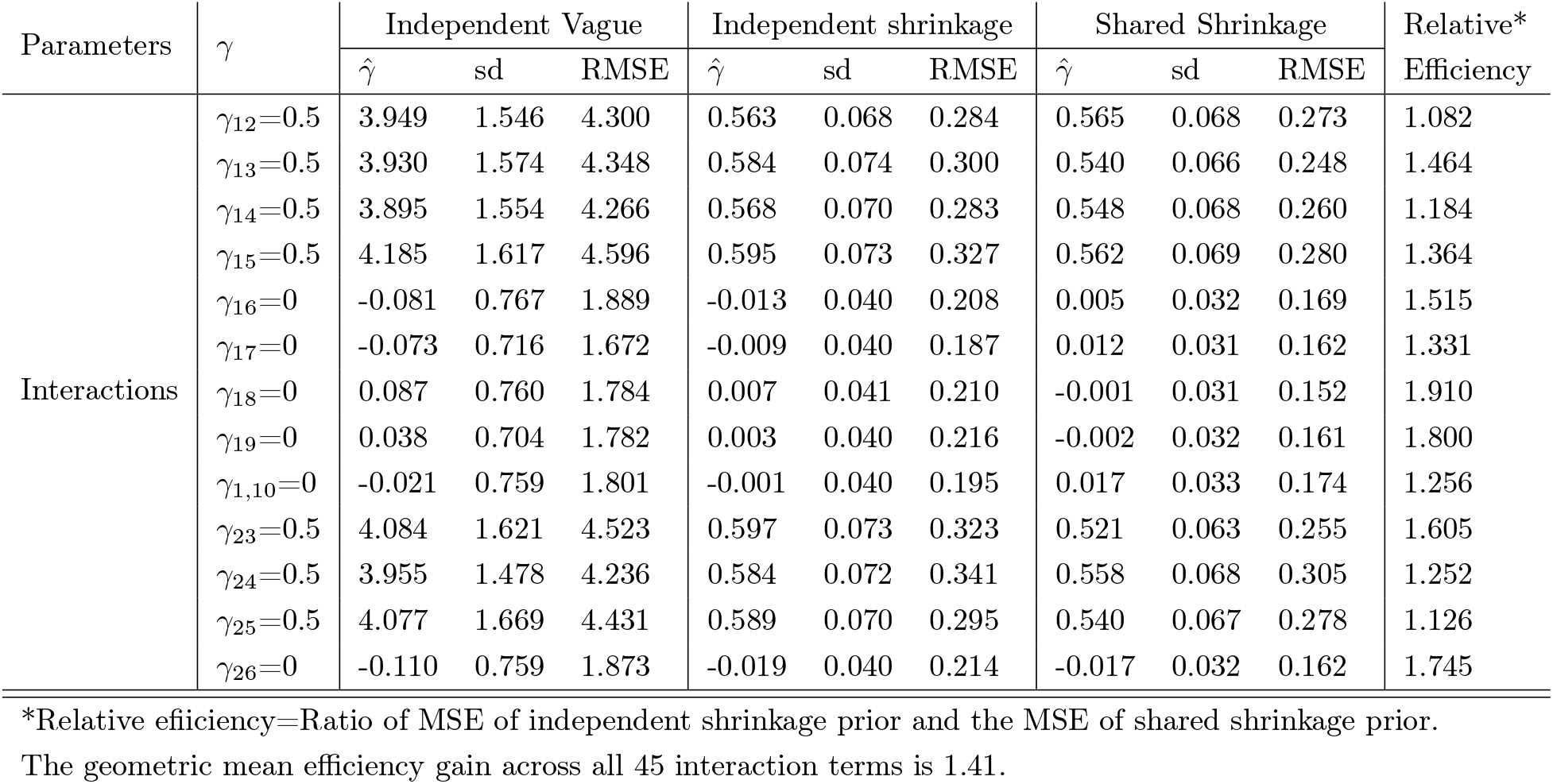
Results from simulation #2: N=250, p=10

| Parameters | $\gamma$ | Independent Vague | | | Independent shrinkage | | | Shared Shrinkage | | | Relative*<br>Efficiency |
| --- | --- | --- | --- | --- | --- | --- | --- | --- | --- | --- | --- |
| | | $\hat{\gamma}$ | sd | RMSE | $\hat{\gamma}$ | sd | RMSE | $\hat{\gamma}$ | sd | RMSE | |
| Interactions | $\gamma_{12}=0.5$ | 3.949 | 1.546 | 4.300 | 0.563 | 0.068 | 0.284 | 0.565 | 0.068 | 0.273 | 1.082 |
| | $\gamma_{13}=0.5$ | 3.930 | 1.574 | 4.348 | 0.584 | 0.074 | 0.300 | 0.540 | 0.066 | 0.248 | 1.464 |
| | $\gamma_{14}=0.5$ | 3.895 | 1.554 | 4.266 | 0.568 | 0.070 | 0.283 | 0.548 | 0.068 | 0.260 | 1.184 |
| | $\gamma_{15}=0.5$ | 4.185 | 1.617 | 4.596 | 0.595 | 0.073 | 0.327 | 0.562 | 0.069 | 0.280 | 1.364 |
| | $\gamma_{16}=0$ | -0.081 | 0.767 | 1.889 | -0.013 | 0.040 | 0.208 | 0.005 | 0.032 | 0.169 | 1.515 |
| | $\gamma_{17}=0$ | -0.073 | 0.716 | 1.672 | -0.009 | 0.040 | 0.187 | 0.012 | 0.031 | 0.162 | 1.331 |
| | $\gamma_{18}=0$ | 0.087 | 0.760 | 1.784 | 0.007 | 0.041 | 0.210 | -0.001 | 0.031 | 0.152 | 1.910 |
| | $\gamma_{19}=0$ | 0.038 | 0.704 | 1.782 | 0.003 | 0.040 | 0.216 | -0.002 | 0.032 | 0.161 | 1.800 |
| | $\gamma_{1,10}=0$ | -0.021 | 0.759 | 1.801 | -0.001 | 0.040 | 0.195 | 0.017 | 0.033 | 0.174 | 1.256 |
| | $\gamma_{23}=0.5$ | 4.084 | 1.621 | 4.523 | 0.597 | 0.073 | 0.323 | 0.521 | 0.063 | 0.255 | 1.605 |
| | $\gamma_{24}=0.5$ | 3.955 | 1.478 | 4.236 | 0.584 | 0.072 | 0.341 | 0.558 | 0.068 | 0.305 | 1.252 |
| | $\gamma_{25}=0.5$ | 4.077 | 1.669 | 4.431 | 0.589 | 0.070 | 0.295 | 0.540 | 0.067 | 0.278 | 1.126 |
| | $\gamma_{26}=0$ | -0.110 | 0.759 | 1.873 | -0.019 | 0.040 | 0.214 | -0.017 | 0.032 | 0.162 | 1.745 |

\*Relative efficiency=Ratio of MSE of independent shrinkage prior and the MSE of shared shrinkage prior.
The geometric mean efficiency gain across all 45 interaction terms is 1.41.

In simulation #3 (Table 3), we evaluated the situation where there were main effects but no interaction effects in the model; this situation is consistent with the shared shrinkage prior specification. Table 4 shows the results for simulation #3 where the average relative efficiency for the shared versus independent shrinkage prior was 2.1. In simulation # 4, we generated data with no main effects but sizeable interaction effects. The results showed that even in the situation that does not directly correspond to the shared shrinkage prior, we found efficiency gains by using this more general prior specification; we found an average efficiency gain of 1.36 across all 45 interaction parameters (geometric mean).

**Table 3:**
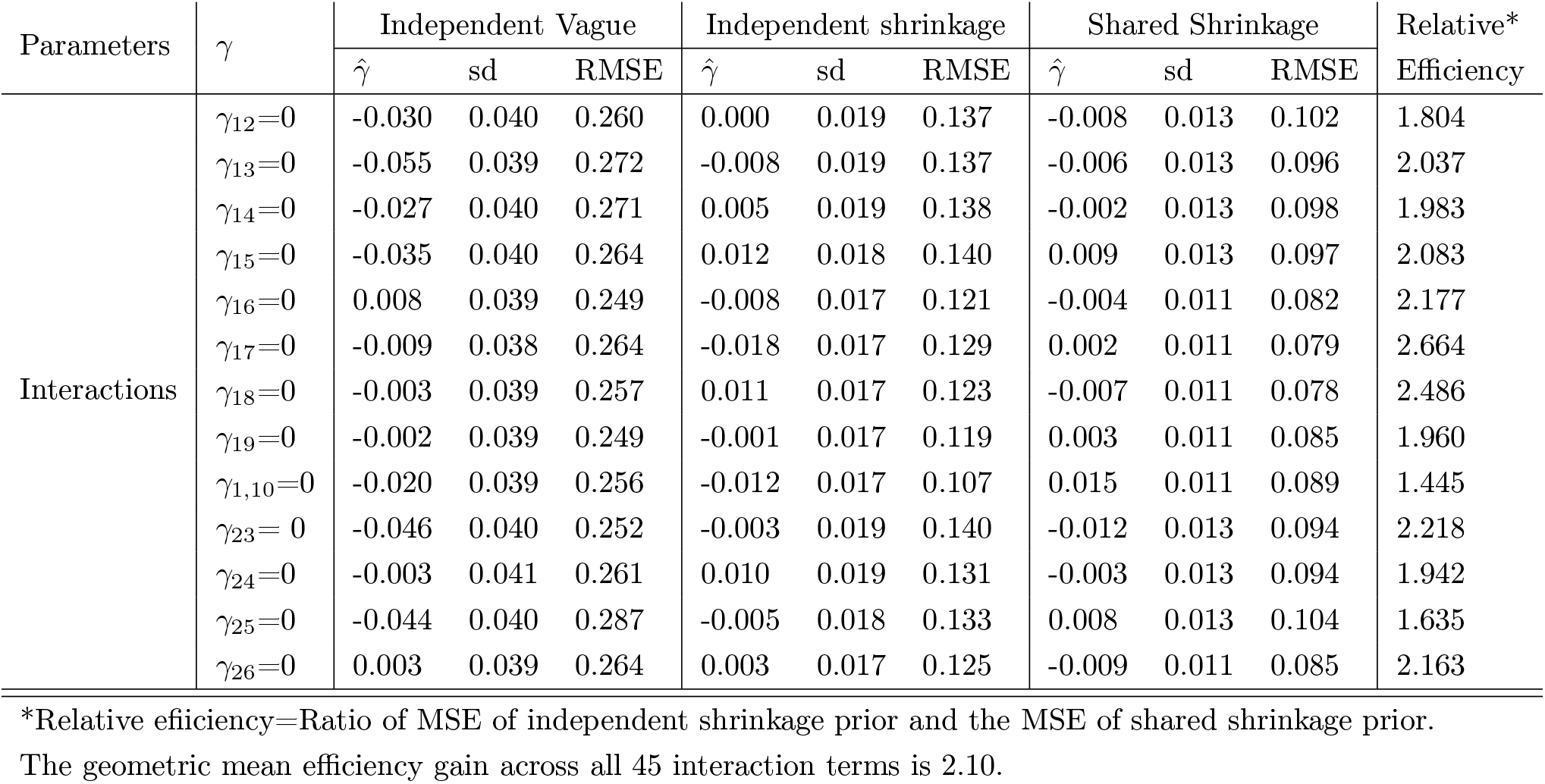
Results from simulation #3: No interaction effects; *N* = 1000, *p* = 10

| Parameters | $\gamma$ | Independent Vague | | | Independent shrinkage | | | Shared Shrinkage | | | Relative*<br>Efficiency |
| --- | --- | --- | --- | --- | --- | --- | --- | --- | --- | --- | --- |
| | | $\hat{\gamma}$ | sd | RMSE | $\hat{\gamma}$ | sd | RMSE | $\hat{\gamma}$ | sd | RMSE | |
| Interactions | $\gamma_{12}=0$ | -0.030 | 0.040 | 0.260 | 0.000 | 0.019 | 0.137 | -0.008 | 0.013 | 0.102 | 1.804 |
| | $\gamma_{13}=0$ | -0.055 | 0.039 | 0.272 | -0.008 | 0.019 | 0.137 | -0.006 | 0.013 | 0.096 | 2.037 |
| | $\gamma_{14}=0$ | -0.027 | 0.040 | 0.271 | 0.005 | 0.019 | 0.138 | -0.002 | 0.013 | 0.098 | 1.983 |
| | $\gamma_{15}=0$ | -0.035 | 0.040 | 0.264 | 0.012 | 0.018 | 0.140 | 0.009 | 0.013 | 0.097 | 2.083 |
| | $\gamma_{16}=0$ | 0.008 | 0.039 | 0.249 | -0.008 | 0.017 | 0.121 | -0.004 | 0.011 | 0.082 | 2.177 |
| | $\gamma_{17}=0$ | -0.009 | 0.038 | 0.264 | -0.018 | 0.017 | 0.129 | 0.002 | 0.011 | 0.079 | 2.664 |
| | $\gamma_{18}=0$ | -0.003 | 0.039 | 0.257 | 0.011 | 0.017 | 0.123 | -0.007 | 0.011 | 0.078 | 2.486 |
| | $\gamma_{19}=0$ | -0.002 | 0.039 | 0.249 | -0.001 | 0.017 | 0.119 | 0.003 | 0.011 | 0.085 | 1.960 |
| | $\gamma_{1,10}=0$ | -0.020 | 0.039 | 0.256 | -0.012 | 0.017 | 0.107 | 0.015 | 0.011 | 0.089 | 1.445 |
| | $\gamma_{23}=0$ | -0.046 | 0.040 | 0.252 | -0.003 | 0.019 | 0.140 | -0.012 | 0.013 | 0.094 | 2.218 |
| | $\gamma_{24}=0$ | -0.003 | 0.041 | 0.261 | 0.010 | 0.019 | 0.131 | -0.003 | 0.013 | 0.094 | 1.942 |
| | $\gamma_{25}=0$ | -0.044 | 0.040 | 0.287 | -0.005 | 0.018 | 0.133 | 0.008 | 0.013 | 0.104 | 1.635 |
| | $\gamma_{26}=0$ | 0.003 | 0.039 | 0.264 | 0.003 | 0.017 | 0.125 | -0.009 | 0.011 | 0.085 | 2.163 |

\*Relative efficiency=Ratio of MSE of independent shrinkage prior and the MSE of shared shrinkage prior.
The geometric mean efficiency gain across all 45 interaction terms is 2.10.

**Table 4:** Results from simulation #4: No main effects; *N* = 1000, *p* = 10

| Parameters | $\gamma$ | Independent Vague | | | Independent shrinkage | | | Shared Shrinkage | | | Relative*<br>Efficiency |
| --- | --- | --- | --- | --- | --- | --- | --- | --- | --- | --- | --- |
| | | $\hat{\gamma}$ | sd | RMSE | $\hat{\gamma}$ | sd | RMSE | $\hat{\gamma}$ | sd | RMSE | |
| Interactions | $\gamma_{12}=0.5$ | 0.745 | 0.021 | 0.307 | 0.579 | 0.015 | 0.154 | 0.559 | 0.015 | 0.138 | 1.245 |
| | $\gamma_{13}=0.5$ | 0.739 | 0.020 | 0.298 | 0.567 | 0.015 | 0.131 | 0.551 | 0.014 | 0.140 | 0.875 |
| | $\gamma_{14}=0.5$ | 0.744 | 0.021 | 0.297 | 0.576 | 0.015 | 0.148 | 0.545 | 0.014 | 0.143 | 1.071 |
| | $\gamma_{15}=0.5$ | 0.752 | 0.021 | 0.308 | 0.594 | 0.015 | 0.162 | 0.557 | 0.015 | 0.135 | 1.440 |
| | $\gamma_{16}=0$ | -0.001 | 0.012 | 0.126 | -0.007 | 0.009 | 0.095 | 0.002 | 0.007 | 0.084 | 1.279 |
| | $\gamma_{17}=0$ | 0.002 | 0.012 | 0.134 | -0.001 | 0.009 | 0.096 | -0.005 | 0.007 | 0.083 | 1.338 |
| | $\gamma_{18}=0$ | 0.000 | 0.012 | 0.139 | 0.004 | 0.009 | 0.095 | 0.007 | 0.007 | 0.077 | 1.522 |
| | $\gamma_{19}=0$ | -0.011 | 0.012 | 0.119 | 0.003 | 0.009 | 0.095 | -0.008 | 0.007 | 0.069 | 1.896 |
| | $\gamma_{1,10}=0$ | 0.000 | 0.012 | 0.139 | -0.009 | 0.009 | 0.100 | -0.006 | 0.007 | 0.074 | 1.826 |
| | $\gamma_{23}=0.5$ | 0.758 | 0.021 | 0.306 | 0.583 | 0.015 | 0.143 | 0.543 | 0.014 | 0.129 | 1.229 |
| | $\gamma_{24}=0.5$ | 0.769 | 0.021 | 0.318 | 0.588 | 0.015 | 0.155 | 0.551 | 0.015 | 0.133 | 1.358 |
| | $\gamma_{25}=0.5$ | 0.748 | 0.020 | 0.296 | 0.570 | 0.015 | 0.149 | 0.548 | 0.014 | 0.125 | 1.421 |
| | $\gamma_{26}=0$ | 0.001 | 0.012 | 0.136 | 0.004 | 0.009 | 0.093 | -0.016 | 0.007 | 0.075 | 1.537 |

\*Relative efficiency=Ratio of MSE of independent shrinkage prior and the MSE of shared shrinkage prior.
The geometric mean efficiency gain across all 45 interaction terms is 1.36.

### 5.2 Simulation for Detection Limit Model

We performed a simulation study for the multi-parameter exposure model that incorporates lower detection limits as described in Section 4. For this case, we considered *p* = 10 main effects and 5 out of these main effects have sizable effect. We need to specify two parameters for each exposure variable: (i) intercept term : *β*_*j*1_*I* (*X*_*ij*_ *≥ C*_*j*_) and (ii) slope term: *β*_*j*2_*I* (*X*_*ij*_ *≥C*_*j*_) (*X*_*ij*_ *−C*_*j*_). For the non-null main effects we consider the effect size *β*_*j*1_ = 0.2 and *β*_*j*1_ = 0.5 for the simulation study. Similarly, for the interaction term of any two main effects we have four parts (i) *γ*_*jk*1_*I* (*X*_*ij*_ *≥C*_*j*_) *I* (*X*_*ik*_ *≥C*_*k*_) (ii) *γ*_*jk*2_(*X*_*ij*_*−C*_*j*_)*I* (*X*_*ij*_ *C*_*j*_) *I* (*X*_*ik*_≥*C*_*k*_) (iii) *γ*_*jk*3_(*X*_*ik*_ *−C*_*k*_)*I* (*X*_*ij*_ *≥C*_*j*_) *I* (*X*_*ik*_*≥ C*_*k*_), and (iv) *γ*_*jk*4_(*X*_*ij*_*− C*_*j*_)(*X*_*ik*_*− C*_*k*_)*I* (*X*_*ij*_ *≥C*_*j*_) *I* (*X*_*ik*_ *≥C*_*k*_). For the interactions terms that have non-null main effects we consider *γ*_*jk*4_ = 0.1 for simulation study. All other *γ*_*jk*1_, *γ*_*jk*2_, and *γ*_*jk*3_’s are set to zero. Similarly as before, we generate the chemical exposure *X*_*ij*_ from a standard normal distribution. For first 5 exposure variables we consider 20% values are below the detection limit and for the rest 10% below the detection limit. Lastly, we consider the overall mean *α* = −2.1 to have a prevalence of 50%. We have a total of 201 parameters in this multi-parameter per exposure model. As the number of parameters are higher than linear model we consider *N* = 1000 observations for simulation study. Table 5 shows the simulation results comparing the shared shrinkage, independent shrinkage and vague prior. Similar to the linear exposure model simulation, we show for this simulation substantial efficiency gain in using shared shrinkage for this more complex model. We also computed the geometric mean efficiency gain of shared shrinkage versus independent shrinkage across all 180 interaction parameters in this multi-parameter model. The mean efficiency gain was 1.37, reflecting substantial gains for the shared shrinkage prior in this sparse data structure.

**Table 5:** Results from simulation for two parameter detection limits model; *N* = 1000, *p* = 10

| Parameters | $\gamma$ | Independent Vague | | | Independent shrinkage | | | Shared Shrinkage | | | Relative*<br>Efficiency |
| --- | --- | --- | --- | --- | --- | --- | --- | --- | --- | --- | --- |
| | | $\hat{\gamma}$ | sd | RMSE | $\hat{\gamma}$ | sd | RMSE | $\hat{\gamma}$ | sd | RMSE | |
| Interactions | $\gamma_{124} = 0.1$ | 0.016 | 0.509 | 0.717 | 0.164 | 0.199 | 0.450 | 0.141 | 0.115 | 0.341 | 1.741 |
| | $\gamma_{134} = 0.1$ | 0.004 | 0.598 | 0.778 | 0.166 | 0.215 | 0.468 | 0.133 | 0.136 | 0.369 | 1.608 |
| | $\gamma_{144} = 0.1$ | -0.039 | 0.577 | 0.770 | 0.111 | 0.204 | 0.451 | 0.116 | 0.146 | 0.382 | 1.394 |
| | $\gamma_{154} = 0.1$ | -0.078 | 0.677 | 0.840 | 0.094 | 0.200 | 0.447 | 0.102 | 0.133 | 0.363 | 1.516 |
| | $\gamma_{164} = 0$ | -0.366 | 3.252 | 1.836 | 0.055 | 0.362 | 0.603 | 0.039 | 0.181 | 0.427 | 1.994 |
| | $\gamma_{174} = 0$ | -0.426 | 2.577 | 1.657 | -0.036 | 0.320 | 0.565 | 0.022 | 0.164 | 0.404 | 1.996 |
| | $\gamma_{184} = 0$ | -0.344 | 1.641 | 1.323 | -0.036 | 0.321 | 0.566 | 0.006 | 0.164 | 0.404 | 1.963 |
| | $\gamma_{194} = 0$ | -0.478 | 1.902 | 1.457 | 0.002 | 0.293 | 0.540 | -0.006 | 0.163 | 0.403 | 1.795 |
| | $\gamma_{264} = 0$ | -0.423 | 2.123 | 1.514 | -0.042 | 0.369 | 0.608 | 0.013 | 0.202 | 0.448 | 1.842 |
| | $\gamma_{274} = 0$ | -0.141 | 0.267 | 0.534 | 0.029 | 0.082 | 0.287 | 0.036 | 0.090 | 0.302 | 0.903 |
| | $\gamma_{284} = 0$ | -0.115 | 0.309 | 0.567 | 0.040 | 0.089 | 0.342 | 0.054 | 0.115 | 0.300 | 1.300 |
| | $\gamma_{294} = 0$ | -0.074 | 0.272 | 0.526 | 0.046 | 0.074 | 0.315 | 0.075 | 0.094 | 0.275 | 1.312 |
| | $\gamma_{344} = 0$ | -0.128 | 0.283 | 0.546 | 0.052 | 0.078 | 0.318 | 0.045 | 0.099 | 0.283 | 1.263 |

\*Relative efficiency=Ratio of MSE of independent shrinkage prior and the MSE of shared shrinkage prior.
The geometric mean efficiency gain across all 180 interaction terms is 1.37.

## 6 NCI-SEER NHL study

The NCI-SEER NHL study (Scosyrev et al., 2012) is a population-based case-control study (508 controls 672 cases) of non-Hodgkin lymphoma (NHL), was designed to determine the associations between exposures of chemicals/pesticides found in used vacuum cleaner bags and the risk of NHL. Different epidemiological pieces of evidence confirm that exposure to chemicals increases the risk of certain cancers in humans(Zahm et al., 1997). Often chemicals enter the household from indoor use or drift-in from outdoor and may persist for months and years in carpet and cushion furniture without being degraded by sunlight, rain, and extreme temperature. Hence carpet dust sampling provides a more objective basis for exposure assessment as it contains integrated pesticide exposure over a long period which is potentially more relevant to disease risk than recent or current exposure. In this study, the samples were collected from used vacuum cleaner bags of 672 cases in Detroit, Iowa, Los Angles, and Seattle and were analyzed for pesticides (Colt et al., 2004). Primarily the laboratory measurements contain missing data due to concentrations being below the minimum detection level. The “fill-in” appproach (Helsel (1990), Colt et al. (2004)) was used for imputation where, for each biomarker, measurements below the detection limit were imputed by first estimating maximum-likelihood estimators of a log-normal distribution from the left-censored data and then imputing values below the detection limit based on this distribution.

Particularly for chemicals with a high percentage of values below their detection limits, results may not be robust to mispecification of the parametric assumptions. The median percent of observations below the detection limit was 61% (across chemicals) with a range of (3% to 93%). Some of the pesticides are highly correlated (> 0.9) and skewed. Figure 1a shows the correlation between the chemicals. In that case, we choose to use only one of them for the analysis and log-transformed the exposure data. In the final data set, we have a total of *p* = 14 chemical exposures on *N* = 1180 individuals (508 controls & 672 cases). Figure 1b shows the correlation between the selected chemicals for the analysis. We considered site, sex, education, and age as covariates (Colt et al., 2004) in all models for our data application. First we estimated a main effect model using a vague prior and an independent shrinkage prior. Figure 2 shows only one chemical, Diazinon, had a 95% HPD interval that excludes zero, for all other chemicals the 95% HPD interval contained zero. Figure 3 shows a random subset of interaction effects estimated under all three prior distributions. The 95% HPD intervals included zero for all estimated interactions for all priors. The intervals for the shared were narrower than the independence shrinkage, and both were narrower than the vague prior.

**Figure 1:**
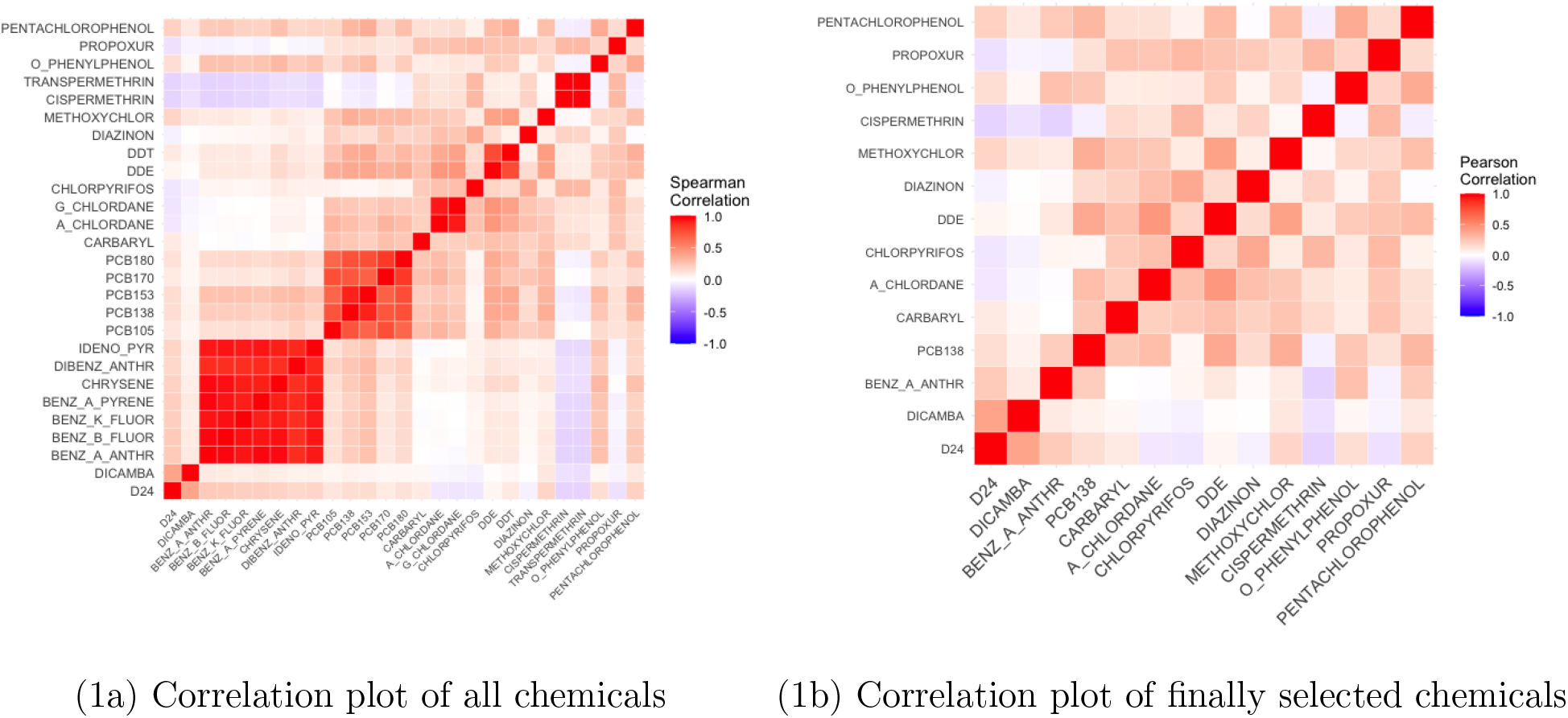
Figure (1a) and Figure (1b) shows the correlation of chemicals.

**Figure 2:**
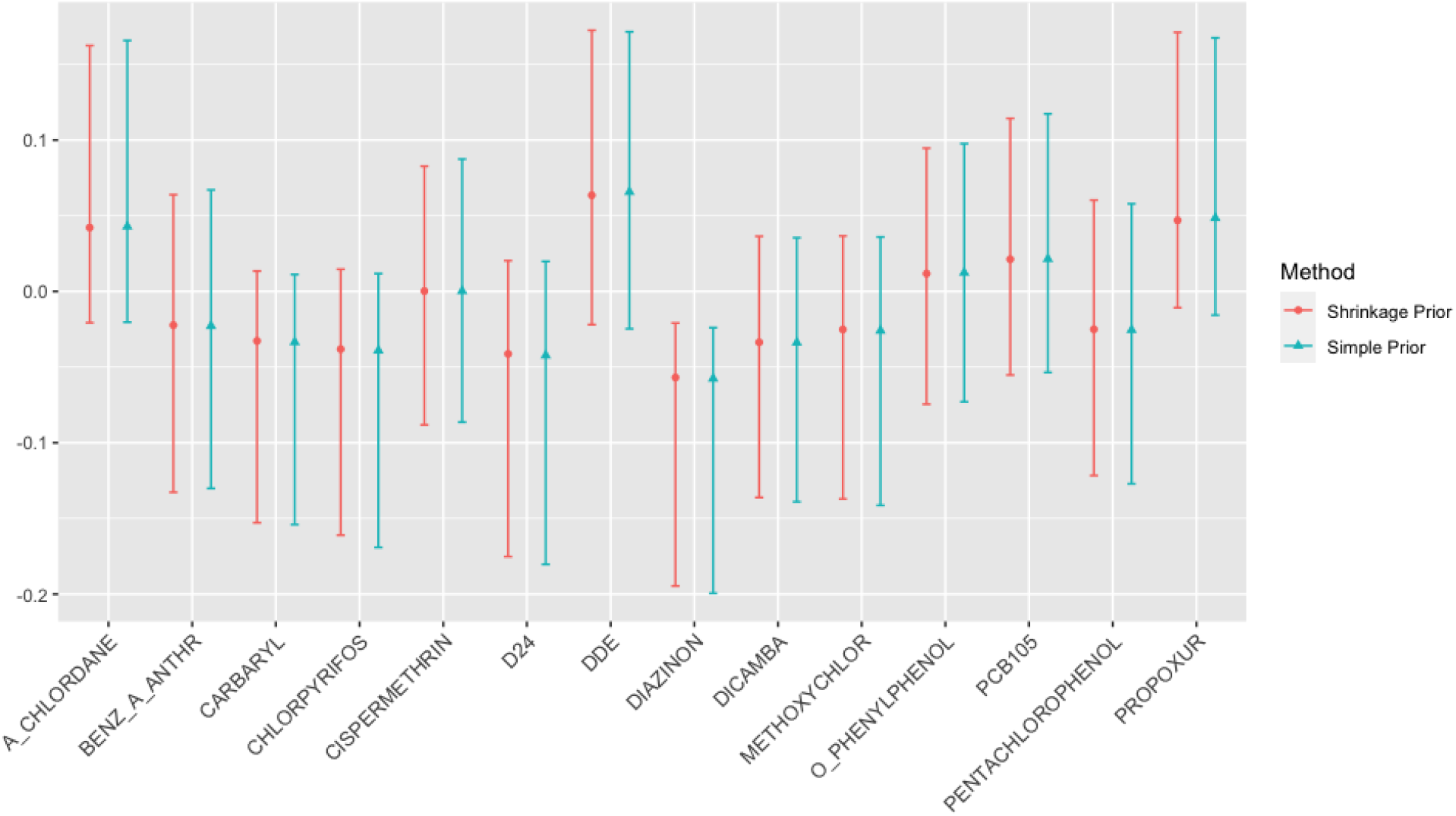
Main effects estimation

**Figure 3:**
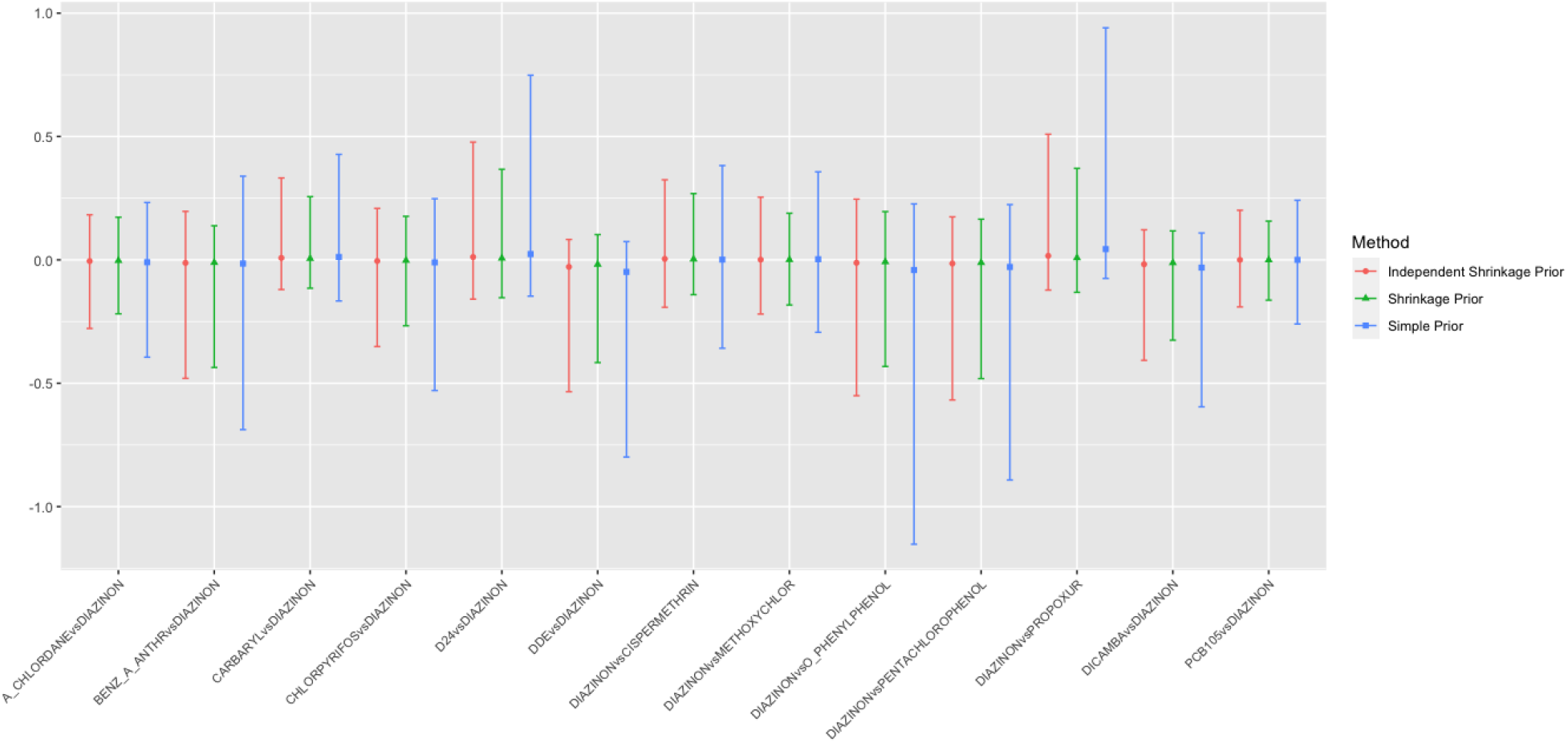
Interaction effect with Diazinon from linear exposure model

The proposed multi-parameter exposure model that incorporates detection limits does not make strong assumptions on the distributions and exposure effects for values below the detection limit. Figures 4 and 5 show the results from fitting the multi-parameter model. These figures show a random set of interaction effects corresponding to the slope-slope interactions (corresponding to *γ*_*jk*4_ parameters). Unlike for the linear exposure models presented, we see strong evidence for interactive effects. Figure 6 shows a contour plot obtained from the estimated terms corresponding to the interaction of D24 and Diazinon. The figure demonstrates a very strong qualitative interaction where exposure effects the risk of NHL only when an individual is exposed to both agents. (note that the log-odds ratios are near zero when either of the agents have exposure at or below their detection limit).

**Figure 4:**
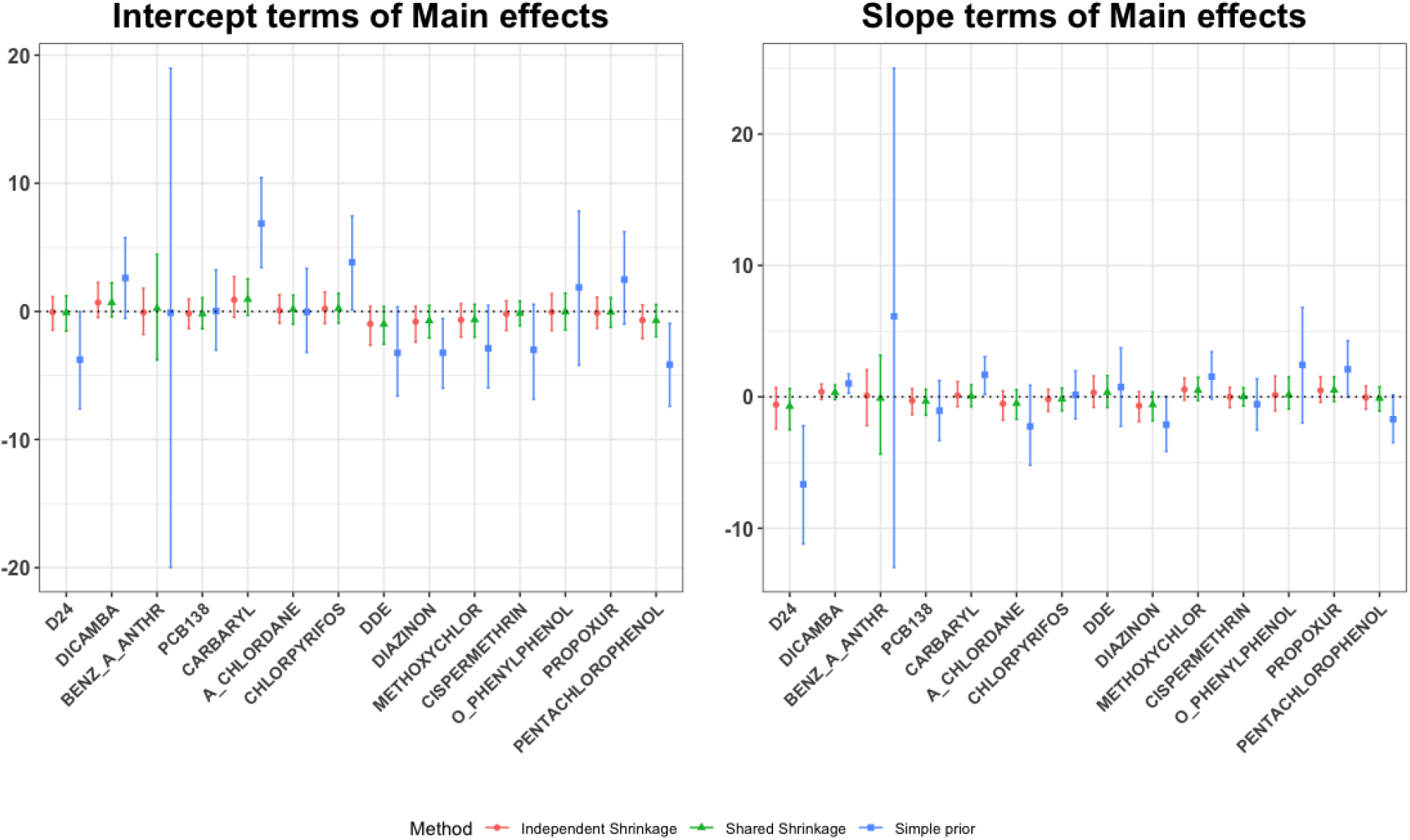
Plots for Intercept terms: *I*(*X*_*ij*_ > *C*_*ij*_) and slopes terms: *I*(*X*_*ij*_ > *C*_*ij*_) (*X*_*ij*_ − *C*_*j*_) for the NCI-SEER NHL data set

**Figure 5:**
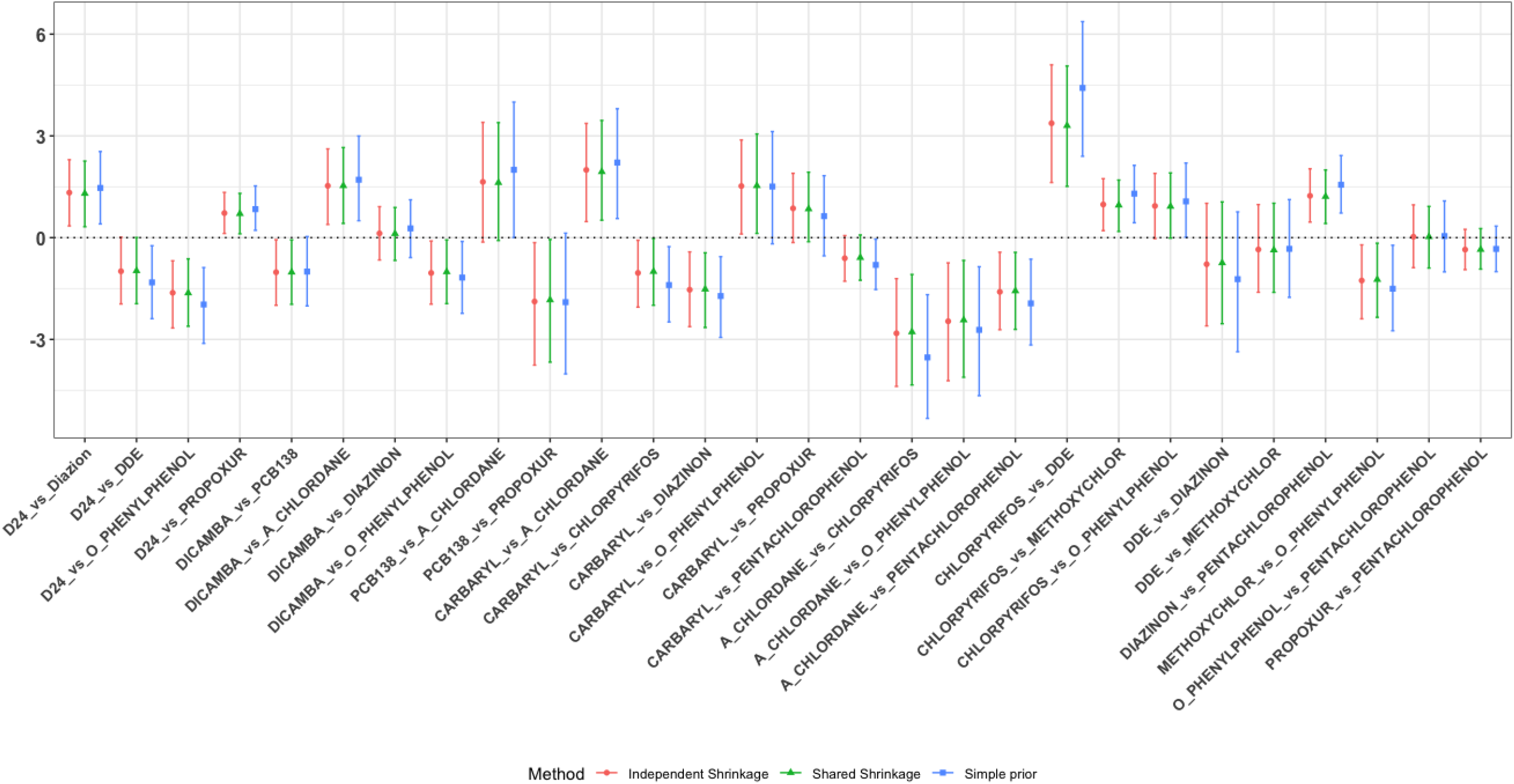
Comparisons between randomly chosen slope vs slope (*γ*_*jk*4_) interaction effects

**Figure 6:**
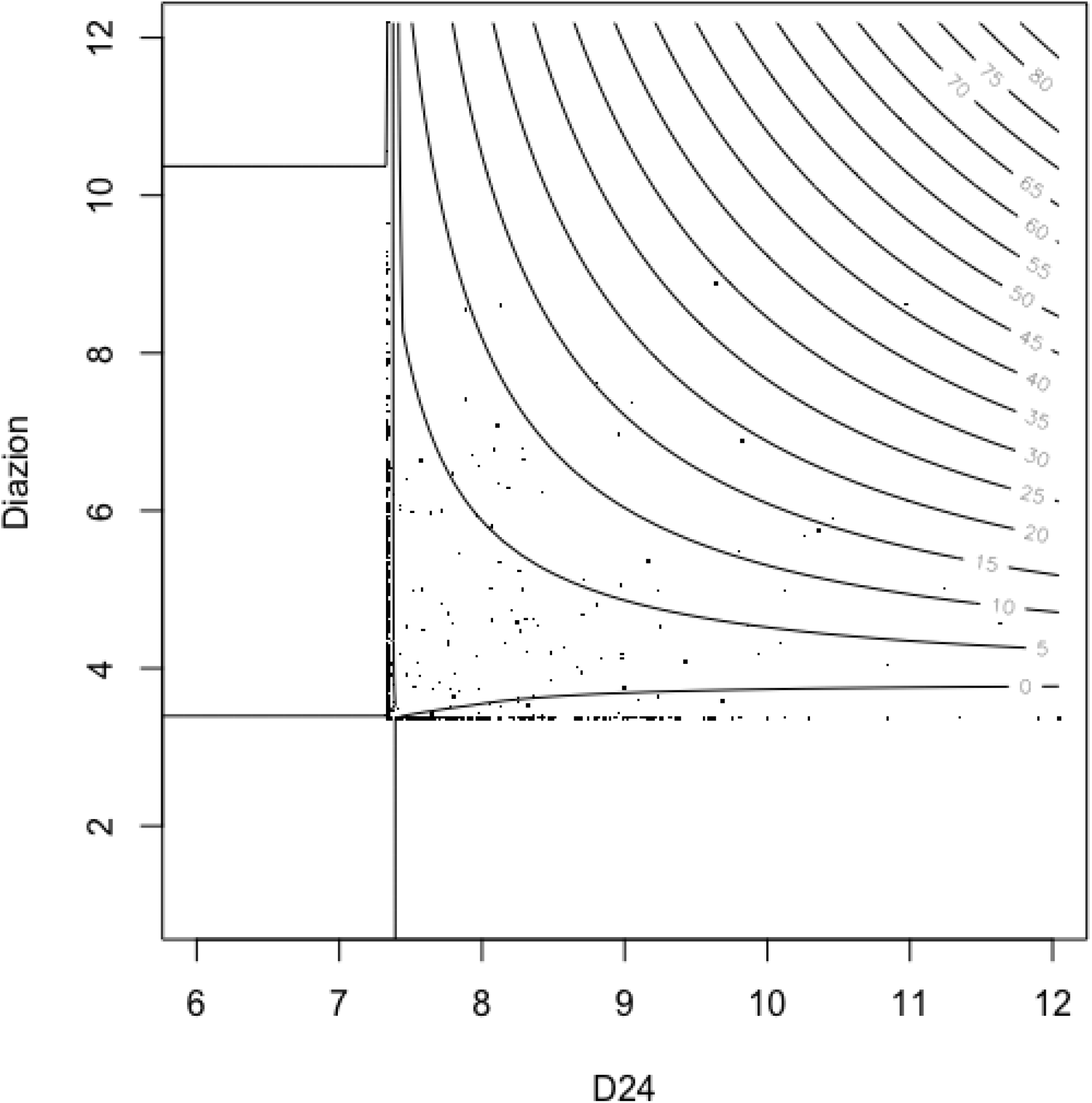
D24 vs Diazion Contour Plot

## 7 Discussion

In this paper we have proposed a shrinkage prior that shares the information between the main effects and interaction effects for estimating the complex relationship between chemical mixtures and disease risk. We proposed a prior that shrinks the interaction term closer to zero when there is a little evidence of a corresponding main effect. This approach is consistent with the hierarchical principle that argues that one should only look for interactions when the corresponding main effects are present. Through simulations, this approach showed sizable efficiency gain in using the shared shrinkage prior when the hierarchical principle holds. Interestingly, we saw that even when there was no main effect, the shared shrinkage prior showed efficiency over priors that did not incorporate this shared information. We presume this is due to the flexibility of the shared shrinkage prior. Bien et al. (2013) proposes a penalized likelihood lasso approach for incorporating the hierarchical principle for parameter estimation. However, there are advantages in using the Bayesian approach over the likelihood approach is that i) there is no need to estimate the penalization constant, and ii) estimating measures of uncertainty are more straightforward using MCMC. Bayesian kernel machine regression (BKMR, Bobb et al. (2015)) is an alternative approach that can be used to identify two-way interactions that does not impose an additive structure in the model formulation. However, this approach does not incorporate the hierarchical principle in the estimation of interaction between mixture components. Further, BKMR cannot be easily extended to flexibly incorporate detection limits as discussed in Section 4.

A major contribution of this work is introducing the shared shrinkage prior in the setting of chemical mixtures. A major analytical issue in the analysis of chemical mixtures is the presence of lower detection limits among the multiple chemicals. We proposed two parameter exposure models that incorporate these detection limits. Specifically for each chemical we include one parameter for the effect of being above the detection limit and another parameter for the linear change (slope) when above this limit. At the expense of adding additional parameters, the proposed approach makes no assumptions about the exposure effects and chemical distributions below detection limits.

The analysis of the NHL mixture data provided interesting insights into the methodology. When using a linear exposure model that imputed values below the detection limits, we found no interactive effects using the independence or shared shrinkage prior. These analysis were based on an imputation approach where chemicals were assumed to be log-normally distributed and values were imputed in the tail of this distribution. The percent of values below the detection limits varied across chemical, but was generally very high (median 61%). We would expect the inferences using imputation and assuming a linear relationship to be highly sensitive to the imposed parametric assumptions. Ortega-Villa et al. (2021) showed the lack of robustness of this type of imputation for a single exposure. When we fit the two parameter exposure model that accounts for detection limits, we found substantially more evidence for interactive effects. Most of these interactions were with regard to D24. For example, we showed very strong positive interactive effect between Diazinon and D24 (see Figure 6) that was missed with the linear exposure with imputation approach.

In the simulations, we considered the case where there were 10 chemicals (*p*=10); the example contained 14 chemicals in the final models. The analysis results were insensitive to the chosen prior distributions. We used a Gamma(1, 1) for the local and global shrinkage priors, but results were very similar when we used either Gamma(1, 0.5) with varaince 0.5 or Gamma(1, 2) with variance 2 distributions. In the simulation studies, with 10 chemicals we have a large number of interactive effects (45 for linear exposure and 180 for two parameter exposure model). With these large number of interactive effects, the simulations and example showed the importance of using shrinkage priors rather than a vague prior that is closely related to maximum-likelihood. For a larger number of chemicals, even the shrinkage prior approaches may have poor performance due to sparsity. In this case, a Bayesian variable selection method (Polson and Scott (2010)) could be used to a smaller number of chemical exposures to include in the model.

We considered models with two-way interactions. Conceptually, we could consider 3 or high order interactions and extend the shared shrinkage idea to this more complex situation. In the spirit of Stone’s generalized additive model Stone (1986), the proposed hierarchical model could be extended to include B-splines rather than polynomial effects. Such an extension would be computational intensive and is left for future work.

## 8 DATA AVAILABILITY STATEMENT

Data will be available upon request. For R code, see the github page.

